## Supplementary File 4 for "CREPE (CREate Primers and Evaluate): a computational tool for large-scale primer design and specificity analysis"

### Overview

CREPE is a batch primer design and specificity analysis tool. It uses Primer3 (https://primer3.org/) to create primers from an input CSV of target sites. It then uses UCSC's In-Silico PCR (https://genome.ucsc.edu/cgi-bin/hgPcr) to identify off-target enrichment sites for each primer pair. Lastly, a custom Python evaluation script (E-script) performs specificity analysis to determine the quality of predicted off-target sites from ISPCR.

The current version of CREPE is designed to create primer pairs for 150bp short-read targeted amplicon sequencing (TAS) applications. As such, it first attempts to find primer pairs that generate a 190bp amplicon that will likely perform well in a TAS experiment. CREPE prioritizes those TAS-optimized (TAS-opt) primer pairs. Through testing, we've found that TAS-opt primer pairs can be made for about 70% of input target sites. For the remaining target sites, CREPE will create alternate primer pairs. This is achieved by giving one of the primers in the reaction a larger area to bind. Relaxed-right primer pairs have the forward primer anchored near the target site, while the reverse primer is given more genomic real estate. Inversely, the Relaxed-left primer pairs have the reverse primer anchored near the target site while the forward primer has a larger region to bind to. Please see the figure below for more detail. The alternate primer amplicons can be as large as 500bp. In this way, CREPE attempts to create functional primer pairs for every input target site.

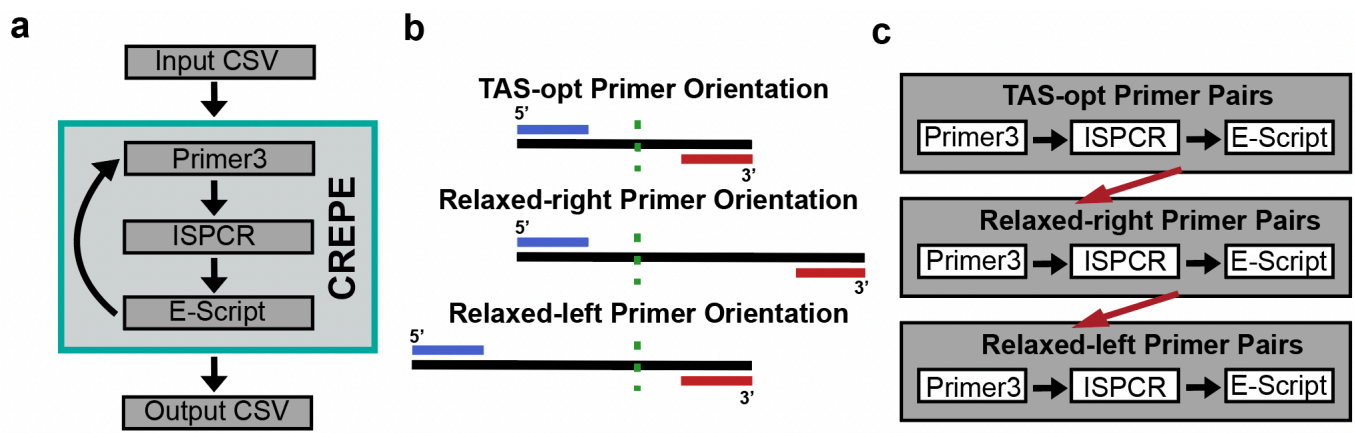

***CREPE requires a custom anaconda environment created during installation. It runs efficiently on local machines, although it can be run on an HPC if that's most convenient.***

### Installation

CREPE runs with few dependencies and a simple conda environment. The suggested installation method is to download and extract the tarball. However, cloning this repository should also work. In any case, the following instructions will help you prepare the files and environment needed for CREPE to run.

#### **1. To begin, download and extract the tar file:**

```
tar -xvzf crepe_download.tar.gz
```

#### **2. Build the CREPE anaconda environment by running the following command:**

```
conda env create --name crepe --file=env.yaml
```

If you have an issue creating the environment from the env.yaml file, then use the following commands to create your environment:

1. `conda create -n crepe python=3.9.12`
2. `conda activate crepe`
3. `conda install pandas=1.4.2`
4. `conda install bioconda::pysam`
5. `conda install conda-forge::biopython=1.79`
6. `conda install bioconda::bedtools`
7. `conda install bioconda::primer3`
8. `conda install bioconda::ispcr`

It should take less than 10 minutes to build the environment.

#### **3. If you have an issue with the tar file, Git clone this directory:**

```
1. git clone -b main https://github.com/martinbreuss/BreussLabPublic
```

### Usage

The input CSV file can contain as many columns as you like. However, the chromosome column must be labeled CHROM, the position must be POS, and the project ID must be PROJ. This is shown in the clinvar10\_demo.csv file.

#### 1. This is the command structure to run CREPE:

```
python CREPE_v1.02.py input.csv ref_genome.fa output_directory projID
```

#### 2. This is an example command:

```
python CREPE_v1.02.py clinvar10_demo.csv hg38.fa test_crepe test_crepe10
```

The final output file with the list of primers is called: FINAL\_OUTPUT\_FILE\_projID (FINAL\_OUTPUT\_FILE\_test\_crepe10). We are working to clean up the output intermediary files. (03/26/2025)

The final output file with all identified off-targets that passed filtering is called: ALL\_OFFTARGETS\_FOUND\_for\_projID (ALL\_OFFTARGETS\_FOUND\_for\_test\_crepe10). Note that this file also contains the on-target primer pair. It's easiest to filter this file to find the off-targets found for specific primers instead of reading the file as a whole.

Below is a sample output from the provided demo file:

| CHROM | POS | PROJ | variant_id | primer3 | isPcr | primer_name | TAS-opt | forward_name | forward_prim | forward_tm | reverse_name | reverse_prim | reverse_tm | amplicon_start | amplicon_end | amplicon_size | primer_count | concerning_off_targets |
| --- | --- | --- | --- | --- | --- | --- | --- | --- | --- | --- | --- | --- | --- | --- | --- | --- | --- | --- |
| 9 | 108912753 | clinvar | clinvar_9_10 | TRUE | TRUE | clinvar_9_10 | FALSE | clinvar_9_10 | CAGTCCAGA | 59.966 | clinvar_9_10 | GGAGATGGA | 59.749 | 108912479 | 108912846 | 367 | 3 | FALSE |
| 7 | 76066259 | clinvar | clinvar_7_76 | TRUE | TRUE | clinvar_7_76 | TRUE | clinvar_7_76 | TAGAGAGAC | 59.818 | clinvar_7_76 | GGATGCCAT | 59.446 | 76066177 | 76066354 | 177 | 1 | FALSE |
| 10 | 43128133 | clinvar | clinvar_10_4 | TRUE | TRUE | clinvar_10_4 | TRUE | clinvar_10_4 | CCAAGGCC | 59.963 | clinvar_10_4 | AGCATCCAG | 57.185 | 43128039 | 43128226 | 187 | 1 | FALSE |
| 1 | 201087921 | clinvar | clinvar_1_20 | TRUE | TRUE | clinvar_1_20 | TRUE | clinvar_1_20 | CAGCACCAC | 59.674 | clinvar_1_20 | CAGGGTTGG | 58.642 | 201087839 | 201088015 | 176 | 1 | FALSE |
| 17 | 61683530 | clinvar | clinvar_17_6 | TRUE | TRUE | clinvar_17_6 | FALSE | clinvar_17_6 | TGCAGAAATC | 57.466 | clinvar_17_6 | AGAAGCCCT | 60.253 | 61683446 | 61683758 | 312 | 1 | FALSE |
| 13 | 33065695 | clinvar | clinvar_13_3 | FALSE | FALSE | NA | NA | check_P | NA | NA | NA | NA | NA | NA | NA | NA | NA | NA |
| 17 | 29055217 | clinvar | clinvar_17_2 | TRUE | TRUE | clinvar_17_2 | TRUE | clinvar_17_2 | GACGTCCAC | 59.898 | clinvar_17_2 | CCAGTGCAA | 60.321 | 29055134 | 29055297 | 163 | 1 | FALSE |
| 14 | 38209841 | clinvar | clinvar_14_3 | TRUE | TRUE | clinvar_14_3 | TRUE | clinvar_14_3 | CTTCGGTGC | 61.485 | clinvar_14_3 | TTACTACCTT | 59.678 | 38209760 | 38209921 | 161 | 3 | TRUE |
| 15 | 42389022 | clinvar | clinvar_15_4 | TRUE | TRUE | clinvar_15_4 | TRUE | clinvar_15_4 | ACGAAGCTC | 59.965 | clinvar_15_4 | CCCACACCI | 59.008 | 42388941 | 42389114 | 173 | 1 | FALSE |
| 20 | 63688197 | clinvar | clinvar_20_6 | TRUE | TRUE | clinvar_20_6 | TRUE | clinvar_20_6 | CTGAGGTCC | 59.404 | clinvar_20_6 | GCAGGGGC | 60.124 | 63688104 | 63688274 | 170 | 2 | FALSE |
| 19 | 50045372 | clinvar | clinvar_19_5 | TRUE | TRUE | clinvar_19_5 | TRUE | clinvar_19_5 | ACCTGTGGC | 60.159 | clinvar_19_5 | TTCTGTGTC | 59.68 | 50045278 | 50045462 | 184 | 2 | TRUE |

### Updates

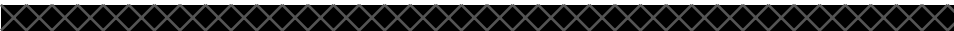

The Python script will be updated to improve performance and allow users to customize amplicon size for applications other than TAS. Please contact Jon, (jonathan[dot]pitsch@cuanschultz[dot]edu), if you have any issues using CREPE.
